# *chronODE*: A framework to integrate time-series multi-omics data based on ordinary differential equations combined with machine learning

**DOI:** 10.1101/2023.12.13.571513

**Authors:** Beatrice Borsari, Mor Frank, Eve S. Wattenberg, Ke Xu, Susanna X. Liu, Xuezhu Yu, Mark Gerstein

## Abstract

Most functional genomic studies are conducted in steady-state conditions, therefore providing a description of molecular processes at a particular moment of cell differentiation or organismal development. Longitudinal studies can offer a deeper understanding of the kinetics underlying epigenetic events and their contribution to defining cell-type-specific transcriptional programs. Here we develop *chronODE*, a mathematical framework based on ordinary differential equations that uniformly models the kinetics of temporal changes in gene expression and chromatin features. *chronODE* employs biologically interpretable parameters that capture tissue-specific kinetics of genes and regulatory elements. We further integrate this framework with a neural-network architecture that can link and predict changes across different data modalities by solving multivariate time-series regressions. Next, we apply this framework to investigate region-specific kinetics of epigenome rewiring in the developing mouse brain, and we demonstrate that changes in chromatin accessibility within regulatory elements can accurately predict changes in the expression of putative target genes over the same time period. Finally, by integrating single-cell ATAC-seq data generated during the same time course, we show that regulatory elements characterized by fast activation kinetics in bulk measurements are active in early-appearing cell types, such as radial glial and other neural progenitors, whereas elements characterized by slow activation kinetics are specific to more differentiated cell types that emerge at later stages of brain development.

## Introduction

Epigenetic mechanisms regulate gene expression and ensure that the information encoded in the genome is correctly translated into cell-type-specific features and functions ^1^. Changes in gene expression naturally modulate cell fate during development and differentiation, and activation or repression of genes at inappropriate times can disrupt normal cellular activity and cause disease ^2–5^. Therefore, establishing the kinetics of gene expression regulation is important for understanding both physiological and pathological processes, and can offer insights on how to artificially switch on or off specific genes for therapeutic purposes ^6–9^.

Most studies so far have investigated the kinetics of RNA production and degradation (i.e., gene expression) ^10–13^, but little is known about the kinetics underlying chromatin changes at regulatory elements and their impact on the expression of target genes. In particular, measurements of chromatin accessibility are considered a proxy for the number of proteins that bind the DNA in a given region of the genome, and among these proteins are transcription factors (TFs) which control the expression of target genes ^14^. As a result, both bulk and single-cell multi-omics experiments have been employed to accurately predict the expression of genes based on the degree of chromatin accessibility at associated regulatory elements ^15–17^. Nevertheless, these predictions are typically constrained to specific steady-state conditions, and cannot be extrapolated to past or future time points in the cell cycle. Deciphering the kinetic parameters of chromatin accessibility changes could potentially allow gene expression to be predicted over a continuous period of time. However, because the timing and rate of chromatin changes are highly context-dependent and vary strongly between genes, computational methods that can precisely model these kinetic parameters at single-gene resolution are needed.

Ordinary differential equations (ODEs) provide an intuitive way to study the kinetics of transcriptional and epigenetic events, since they model the rate of change of a dependent variable (i.e., RNA molecules or chromatin accessibility) with respect to an independent variable (i.e., time). In recent years, several studies have applied ODE frameworks to infer the trajectory (i.e., velocity) of cells in future time points, based on the collective directions and rates of transcriptional and epigenetic changes of genes ^18–21^. These methods typically leverage a one-time, single-cell snapshot taken during a dynamic process to capture cells at different stages, and construct a latent time which describes the temporal progression of cells without being constrained by actual time-series measurements. This latent time is then used to estimate the kinetic parameters and switch times for the activation of individual genes, based on their level of chromatin accessibility or ratio between spliced and unspliced mRNA. Finally, the velocities of all genes in a given cell are combined to predict the trajectory of the cell over time. While this is a valid strategy to overcome the lack of technologies that can monitor the same single cell over time, it poses some limitations for accurately estimating kinetic parameters at the resolution of individual genes. First, these methods employ theoretical ODEs that are not directly formulated from real data, hence their biological interpretation of the temporal kinetics may be limited. Second, kinetic parameters are estimated from a latent time, and therefore lack validation from time-resolved measurements.

In comparison to velocity methods applied to one-time snapshot measurements, modeling real time-series genomic data is a more suitable way to formulate kinetic equations that can accurately describe dynamics of gene expression and chromatin accessibility over time. Here, we develop *chronODE*, a framework based on ODEs to precisely quantify the rate of change in functional genomic signals over time. Using this modeling approach we can formulate ODEs in ways in which their kinetic parameters have a clear biological meaning with respect to time. By integrating bulk and single-cell chromatin accessibility data generated at eight time points during mouse brain development ^22,23^, we identify candidate *cis*-regulatory elements (cCREs) with diverging time-series patterns that recapitulate their activity in specific cell types. Finally, we apply this modeling framework to RNA-seq data generated during the same time course ^24^ and demonstrate that the rate of change in chromatin accessibility at cCREs can accurately predict the rate of expression changes at target genes over time.

## Results

### Rewiring of the accessible genome during brain development shows region-specific timing

We analyzed time-series maps of chromatin accessibility generated for three regions of the mouse brain at eight developmental time points (forebrain, midbrain and hindbrain; from day E10.5 to the first postnatal day; Supplementary Table 1) ^22^. We designed a protocol for data normalization and batch correction that allowed us to integrate DNase-seq and ATAC-seq data from the same region in one single time course (Supplementary Figure 1). Starting from the ENCODE registry of mouse cCREs (*n* = 926,843) ^25^, we identified 405,554 cCREs with signatures of active chromatin in at least one region and time point. On average across the three regions, 24% of active cCREs showed dynamic changes in chromatin accessibility over time, with the largest number observed in the forebrain (*n* = 148,908; 37%; Figure 1a).

**Figure 1.**
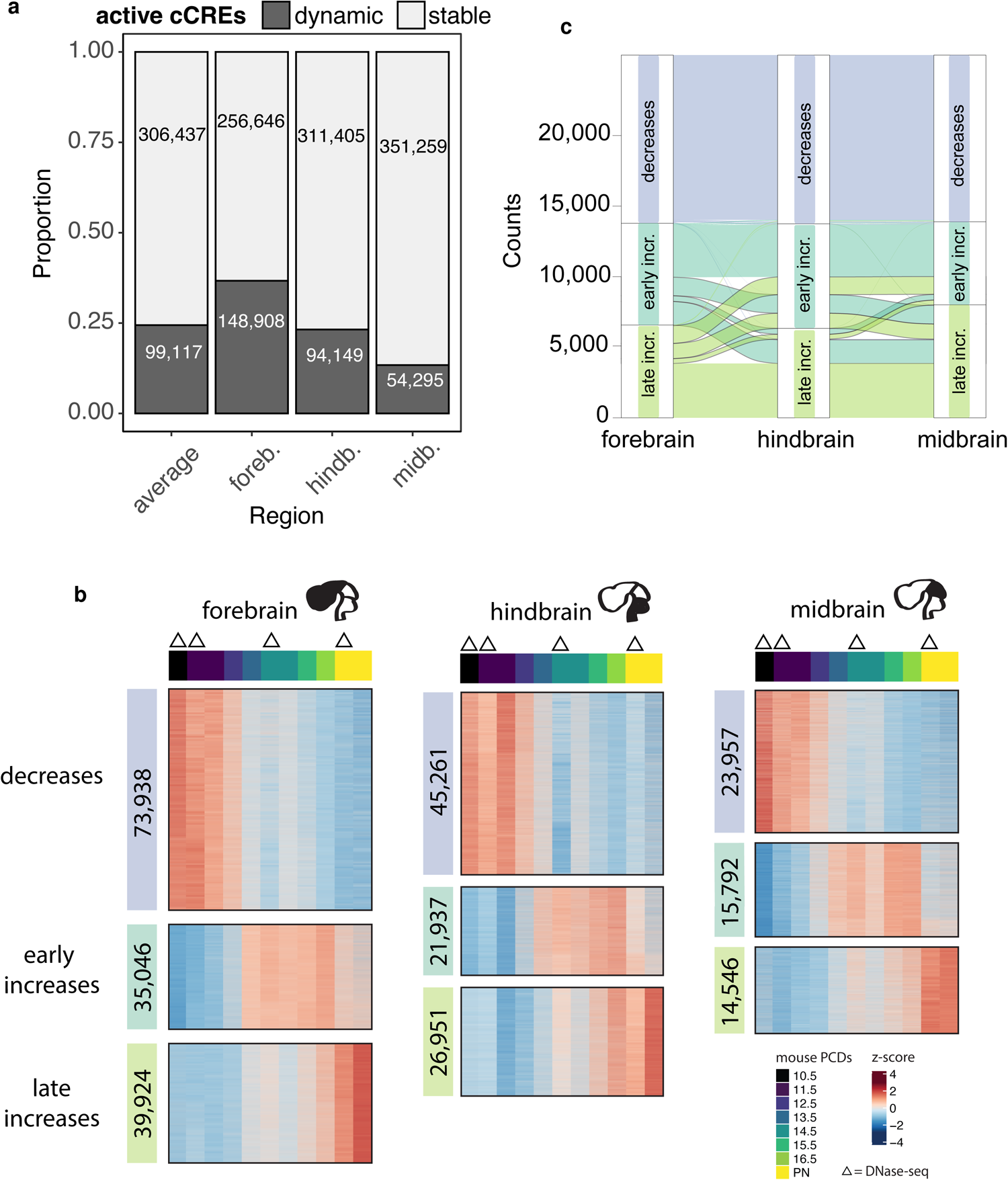
Dynamic candidate cis-regulatory elements (cCREs) during brain development show region-specific timing. **a**: Barplot showing the proportion (y axis) of active cCREs that show dynamic changes of chromatin accessibility during the time course. We show the proportion for each region (forebrain, midbrain, hindbrain) as well as the average across the three regions (x axis). **b**: Chromatin accessibility profiles of dynamic cCREs (rows) across the eight time points (columns) can be grouped into three main clusters: decreasing, early increasing, and late increasing. The clusters are color-coded and the numbers of cCREs in each cluster are indicated. The upper colored legend indicates the time points. For time-points E11.5, E14.5 and PN we display the signal obtained from both ATAC-seq and DNase-seq maps. Signals corresponding to DNase-seq maps are indicated by a triangle. The profiles consist of row-normalized z-scores. PN: first post-natal day. **c**: Alluvial plot showing the proportion of dynamic cCREs (y axis) that show concordant and discordant patterns across the three regions (x axis). In this case we considered a subset of 25,703 cCREs that are dynamic in all three regions.

Roughly similar proportions of cCREs showed increasing and decreasing profiles of chromatin accessibility (53% and 47% on average across the three regions, respectively). Most decreasing patterns were constrained to an early developmental window (E12.5-13.5) and were largely conserved across the three regions, suggesting that these changes may involve a general coordinated transition from pluripotency to more specialized cell types (Figure 1b-c). In contrast, increasing patterns were more varied. Increases could happen either early or late in development (mostly between E12.5-E13.5 or postnatally, respectively; Figure 1b). This timing also varied strongly between regions for the same cCRE. For instance, nearly 49% of early-increasing cCREs in the forebrain were classified as late-increasing in at least one of the other two regions (Figure 1c and Supplementary Table 2). Albeit qualitative, this first classification indicates that not all cCREs undergo chromatin changes at the same time and rate. Additionally, the activation and inactivation kinetics of the same cCRE may vary among different regions, and we hypothesized that these variations could be associated with the cellular context specific to each brain region.

### *chronODE* infers time-series trajectories and highlights cell type-specific kinetics of chromatin accessibility

To investigate the potential context-specific activation rates of cCREs with a more quantitative approach, we have developed *chronODE*, a time-series framework based on ordinary differential equations (ODEs). In this framework, we define the degree of chromatin accessibility at a given cCRE as a time-dependent variable *y*. We assume that *y* will reach saturation (steady-state) at a given point in time, although some cCREs may not reach this saturation point during the window of time monitored in the current study. Practically, this means that the maximum degree of chromatin accessibility at a given cCRE approaches a limit, which in single-cell experiments is proportional to the number of nucleosomes that overlap the cCRE, and in the case of bulk experiments, it is also proportional to the total number of sequenced cells.

In this context, the rate of change in chromatin accessibility at a given time point *t* can be defined by a non-linear function *f*, that is

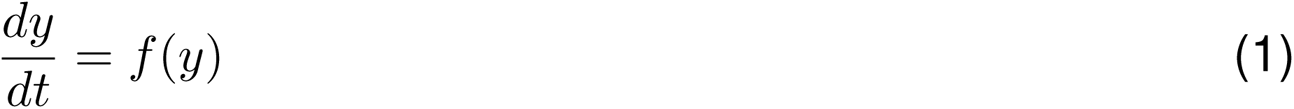

Under this scenario, the function describing how *y* varies over time can be theoretically approximated by a logistic curve, which has been previously employed to model growth and decay population dynamics ^26,27^, and we propose the following first-order differential equation addressing an initial value problem:

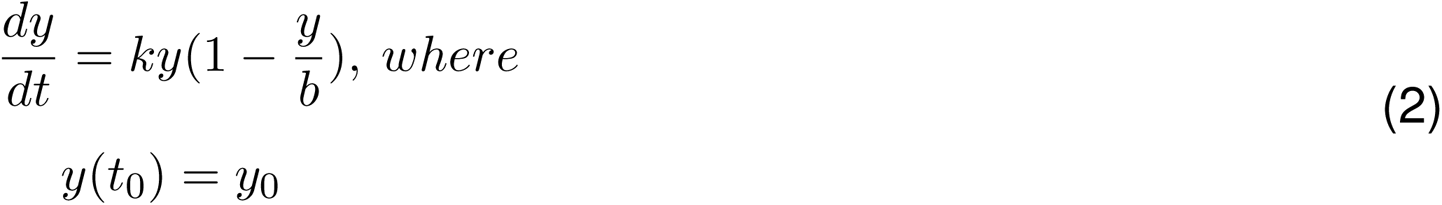

When considering a logistic increasing or decreasing curve (Figure 2a-b), *k* represents the rate of change in chromatin accessibility and *b* represents the horizontal asymptote of *y*. However, this formula can also mathematically accommodate time-series profiles that match only portions of the logistic curve, such as log-like or exponential patterns (Figure 2c-f). Thanks to the flexibility of this framework we could fit this ODE to model the time-series profiles of *>*99% of our dynamic cCREs.

**Figure 2:**
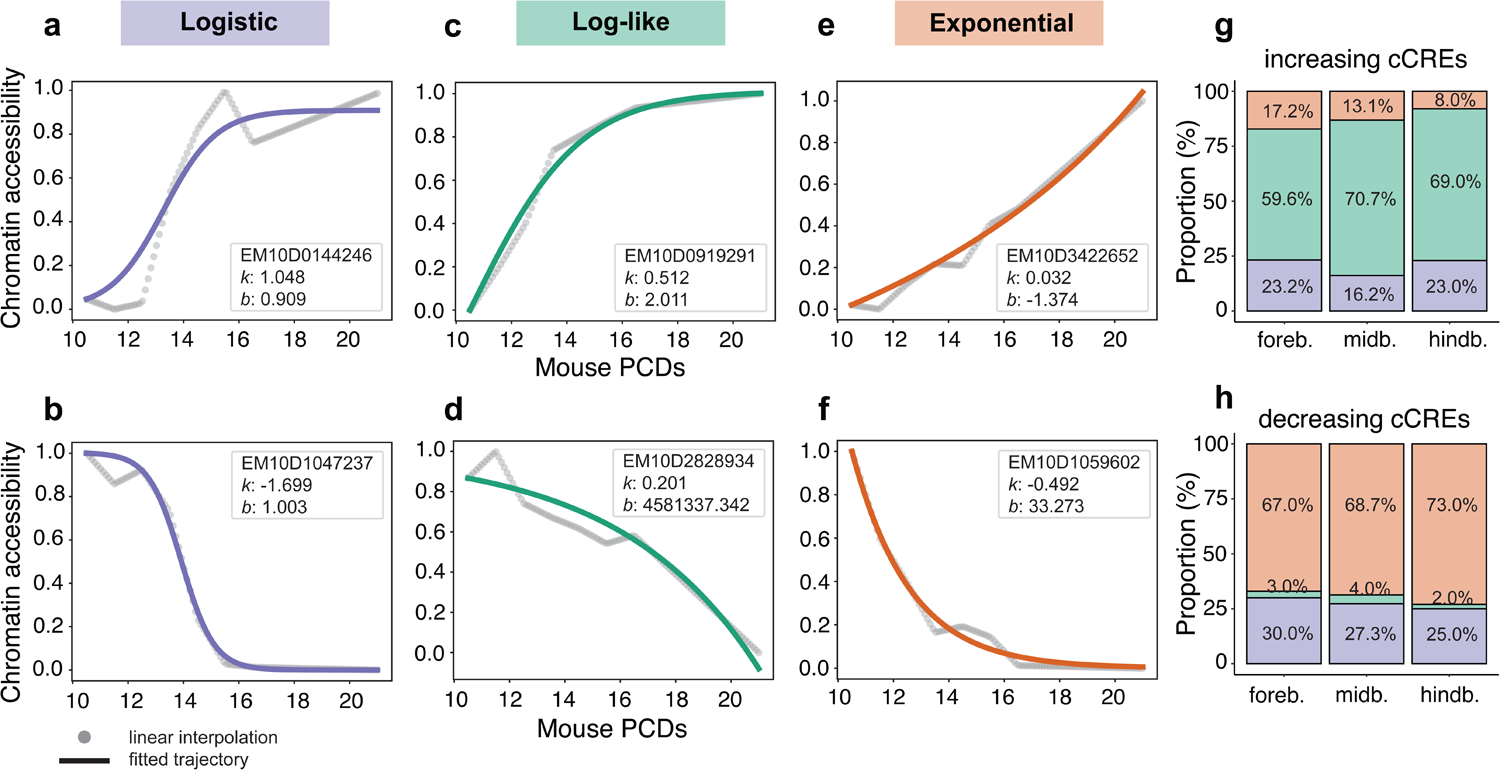
Main signal trajectories modeled by chronODE. **a-f:** Examples of chromatin accessibility trajectories following the logistic curve (a-b), and the log-like (c-d) and exponential (e-f) portions of it. For example we display the cCRE ENCODE identifier with the corresponding and *b* parameters. For each example we display the cCRE ENCODE identifier with the corresponding *k* and *b* parameters. **g-h**: Barplots showing for each brain region (x axis) the proportion of cCREs (y axis) characterized by logistic, log-like and exponential patterns. Increasing cCREs: upper panel; decreasing cCREs: lower panel.

This modeling approach offers a number of advantages. First, it employs real time-series data to infer the kinetic parameters, differently from previous methods that rely on a latent time to fit the ODE solution ^18–21^. Second, it offers greater intuitiveness compared to fitting a high-order polynomial function, as the kinetics are defined by only two parameters, which are biologically interpretable. Third, it can be expanded to model, under similar assumptions, time-series changes in gene expression or other chromatin features such as histone modifications.

We numerically solved equation (2) to fit the kinetic parameters *k* and *b* for every dynamic cCRE across the three brain regions and reconstruct the most likely time-series trajectory (Supplementary Figure 2a). We then classified the trajectory of each cCRE as either logistic, log-like or exponential patterns (Supplementary Figure 2b-c). Most increasing cCREs followed either log-like or logistic patterns, whereas only a minor fraction showed slow, exponential increases (Figure 2g and Supplementary Figure 3a). Instead, most decreasing cCREs exhibited exponential or logistic patterns, and almost never displayed slow, log-like curves (Figure 2h and Supplementary Figure 3a).

Consistent with our initial observations, we found that early-increasing cCREs showed higher *|k|* values compared to late-increasing cCREs (Figure 1b and Supplementary Figure 3b). Comparing, for a given cCRE, the magnitude of *|k|* across the three regions reveals more granular changes in the kinetics of chromatin accessibility than our initial exploratory analyses had suggested. For instance, even when considering cCREs that were systematically early- or late-increasing in all three regions we could identify some with a much faster activation rate in one specific region (Figure 3a). This was particularly evident in the case of decreasing cCREs, for which our initial clustering analysis did not highlight strong variations in the timing of chromatin closing (Figures 3b and 1b).

**Figure 3:**
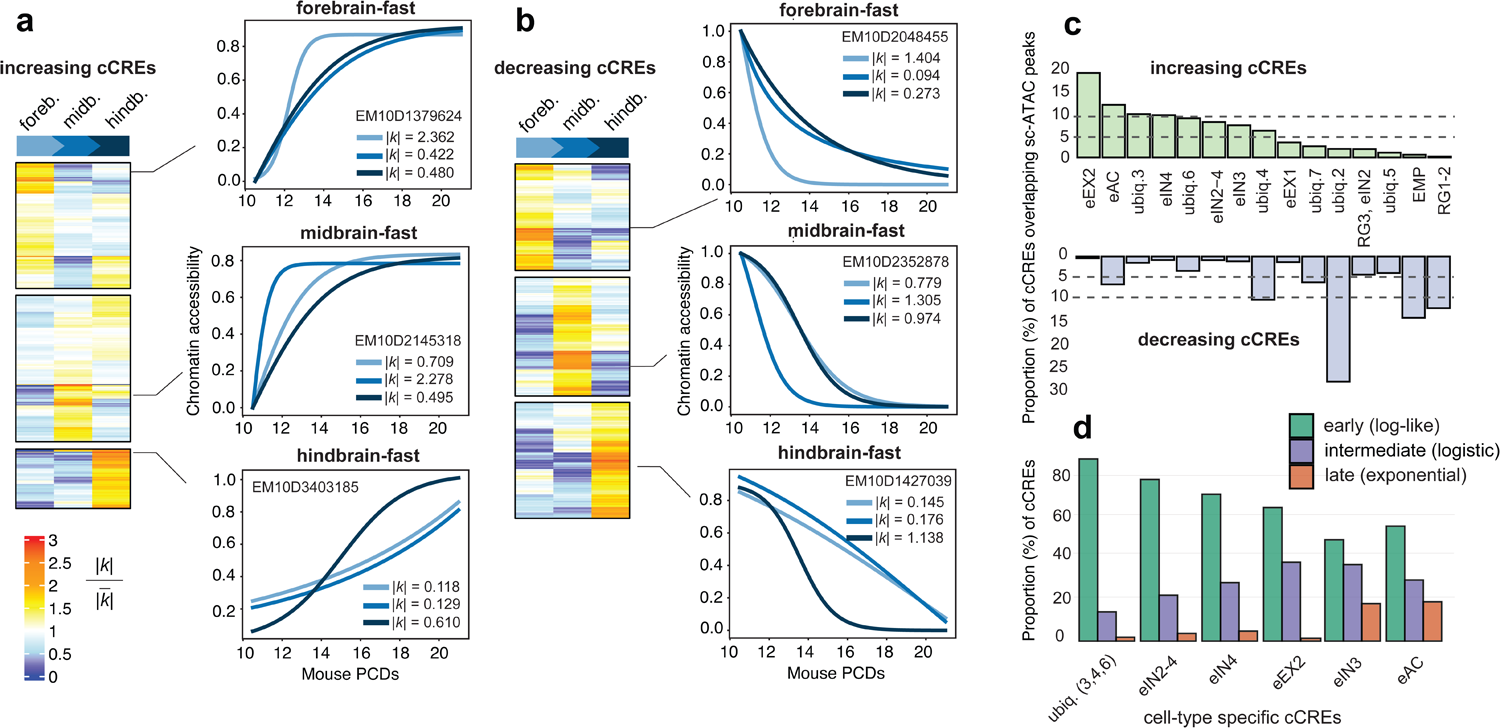
chronODE provides biologically interpretable kinetic parameters. **a-b:** Heatmaps showing for increasing and decreasing cCREs (rows) changes in the kinetic parameter across the three brain regions (columns). To facilitate the comparison, we display a normalized value of (i.e., | | divided by the average | | across the three regions) and consider a subset of cCREs that are characterized by the same pattern (decreasing, late increasing, early increasing) in all three regions. Zoom-in lineplots show examples of cCREs characterized by faster kinetics in a particular region;for each example we report the cCRE ID and the corresponding value of |k| in each region. **c:** Barplot showing the proportion (%) of forebrain increasing and decreasing cCREs (y axis) that overlap cell-type specific peaks identified by scATAC-seq experiments (Preissl et al., 2018; see Supplementary Table 3). Horizontal dashed lines indicate 5% and 10% percentages. **d:** Barplot showing the proportion (%) of forebrain increasing cCREs (y axis) characterized by log-like (green), logistic (purple) and exponential (orange) trajectories across different cell-type specific sets (x axis). Only sets showing a proportion >5% in panel care displayed. We merged ubiquitous-3, −4, and −6 sets into one set. intermediate and late patterns.

These regional kinetic differences align with the hypothesis that the rate of activation and inactivation of regulatory elements may strongly depend on the cellular composition of each brain region. Following this assumption, ubiquitously active cCREs may exhibit very different kinetics from cell-type specific cCREs, whose activation rates may be influenced by the changing regional abundance of the specific cell type over time. To explore this possibility, we analyzed single-cell (sc) ATAC-seq data generated in the forebrain during the same time course as the bulk regional data (i.e., from E11.5 through the first PN day) ^23^. Previous analyses of these data highlighted the temporally regulated appearance of differentiated cell types, in particular mature excitatory neurons (eEX2, between E12.5 and E13.5), mature inhibitory neurons (eIN4, between E12.5 and E13.5, and eIN3 after E14.5), and astrocytes (eAC, after E16.5) ^23^. Consistent with these previous observations, our forebrain-increasing cCREs showed the largest overlap with peaks specific to eEX2, eAC and eIN4, and the least overlap with peaks specific to neuronal progenitors (radial glia, RG1-2) and erythromyeloid progenitors (EMP) (Figure 3c). The latter were instead particularly abundant among our set of forebrain-decreasing cCREs, consistent with EMP and RG1-2 progressively disappearing during the time course ^23^ (Figure 3c).

We expected that increasing cCREs specific to late-emerging cell types would show different types of trajectories compared to ubiquitous cCREs, the majority of which are active in early-emerging cell types such as radial glial and other neuronal progenitors (Supplementary Table 3). Indeed, we found that the proportions of early (log-like), intermediate (logistic) and late (exponential) trajectories were inversely correlated across the different sets of cCREs, consistent with the order of temporal appearance of the corresponding cell types (Figure 3d). Compared to ubiquitous cCREs, which showed the largest proportion of early trajectories, those cCREs specific to eEX2 and eIN4 (appearing between E12.5 and E13.5) displayed higher proportions of intermediate trajectories. Finally, cCREs specific to the late-emerging eIN3 and eAC populations (appearing after E14.5 and E16.5, respectively) reported the largest proportions of late patterns. Altogether, these results suggest that trajectories of chromatin accessibility inferred from bulk experiments can recapitulate the emergence of cell types during development, wherein cCREs specific to earlier cell types exhibit early patterns, while cCREs specific to later cell types are characterized by

### Genes linked to multi-pattern cCREs are more dynamic during brain development and are enriched in brain-specific functions

Having established a framework for modeling changes in chromatin accessibility at regulatory elements, we next sought to investigate how these changes can impact the expression of target genes over time. To achieve this goal, we compiled a catalog of cCRE-gene pairs, by linking each cCRE to its nearest protein-coding gene based on linear distance ^28,29^. Employing this proximity-based method, putative target genes were linked to multiple cCREs, consistent with prior findings ^29^. Notably, a considerable fraction (on average 27%) of these genes exhibited not only quantitative but also qualitative multi-assignment, i.e. they were linked to cCREs displaying all three major time-series patterns (decreasing, early-increasing, or late-increasing; multi-pattern regulated genes; Figure 4a). 46% of the genes were associated with just one type of cCREs (mono-pattern regulated genes).

**Figure 4:**
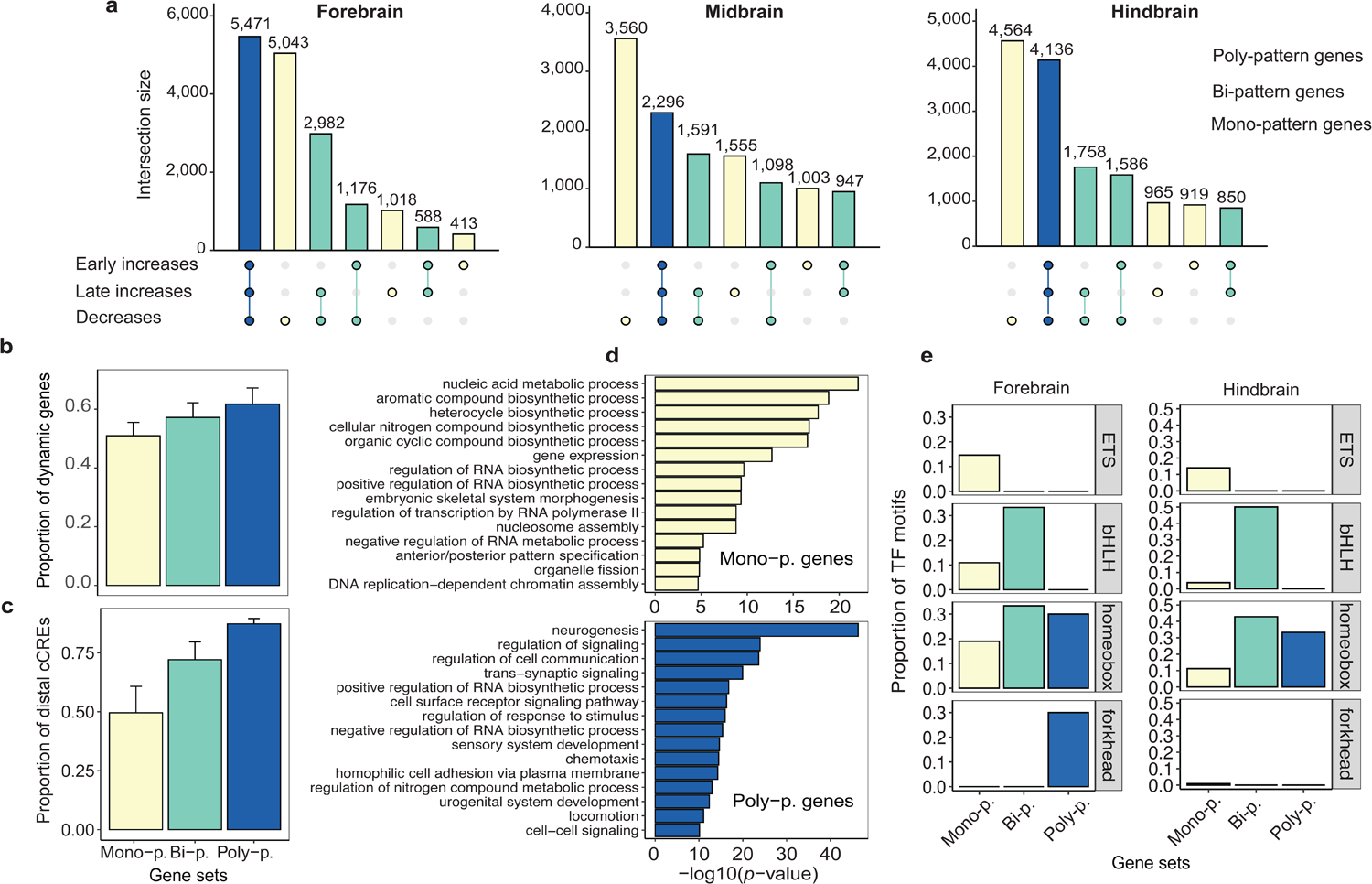
Genes linked to multi-pattern cCREs are more dynamic during brain development and are enriched in brain-specific functions. **a:** Upset plots showing, for each brain region, the number of genes (y axis) that are associated with decreasing, early increasing and late increasing cCREs. Mono-, bi- and poly-pattern genes are associated with one, two, and three types of cCREs, respectively (light yellow / green / blue). **b:** Barplot showing the proportion of dynamic genes (y axis) across sets of mono-, bi-, and poly-pattern genes (x axis). Dynamic genes are defined as those showing significant changes in gene expression over time (maSigPro FDR < 0.01). We report mean proportion and standard deviation across the three regions. **c:** Barplot showing the proportion of distal cCREs (y axis) across sets of mono-, bi-, and poly-pattern genes (x axis). Distal cCREs are defined as those located > ±2 Kb from an annotated transcription start site. We report mean proportion and standard deviation across the three regions. d: Barplot showing Gene Ontology terms for biological process (y axis) enriched among mono- and poly-pattern genes with the corresponding -log10 p-values (x axis). **e:** Barplot showing the proportion of motifs from ETS (Erythroblast Transformation Specific), bHLH (basic Helix-Loop-Helix), homeobox, and forkhead TF families (y-axis) enriched at the promoters of mono-, bi-, and poly-pattern genes in forebrain (left) and hindbrain (right).

We found that the variety of regulatory patterns associated with a gene correlates with several gene features. Consistently across the three brain regions, multi-pattern genes were paired with more distal cCREs and displayed more significant changes in expression over time compared to mono-pattern genes (Figure 4b-c). In line with these results, which suggest a more prominent role of multi-pattern genes during brain development, we found them to be enriched in neural and brain-specific functions, including neurogenesis, trans-synaptic signaling, and sensory system development (Figure 4d). Multi-pattern genes also showed enrichment of motifs corresponding to homeobox and forkhead TFs, which have been reported to orchestrate key processes during brain development, such as regional specification, neuronal differentiation and axonal guidance and connectivity ^30–36^ (Figure 4e). In contrast, mono-pattern genes were enriched in housekeeping functions such as gene expression and compound biosynthesis, and displayed modest motif enrichment also for other TF families (e.g., ETS and bHLH; Figure 4d-e).

Altogether, these results suggest that the precise expression of genes essential for brain development may be governed by a more complex regulatory network. This regulatory control appears to involve cCREs with diverse time-series trajectories, highlighting a sophisticated orchestration of essential genetic processes. In contrast, less critical genes or those associated with non-brain specific functions may rely on a simpler regulome, potentially reflecting a differential degree of control based on the biological significance and impact of these genes.

### Predicting time-series gene expression patterns from changes in associated cCRE activity

Besides revealing a potential correlation among the diversity of regulatory patterns linked to a gene, its TF regulatory network and its functional role during brain development, these results also suggest that accurately predicting changes in a gene’s expression levels during development based on changes in chromatin accessibility within its cCREs may require either a single- or multi-cCRE schema model.

Here, in order to assess whether changes in gene expression can be predicted by chromatin changes at the corresponding cCREs (e.g., chromatin accessibility), we considered the simple single-cCRE scenario. Thus, given a cCRE-gene pair that are linked only to each other, we employed the ODE-inferred time-series derivatives of chromatin accessibility of the cCRE as input to predict the time-series derivatives of expression of the gene. Specifically, we solve a multivariate time-series regression where the change over time *t* in the expression of gene *g_n_* depends on changes over time *t* in all the chromatin features *c_i_*associated with the gene (note that here we employ only the chromatin accessibility feature and one cCRE per gene, thus *i* = 0) following this equation:

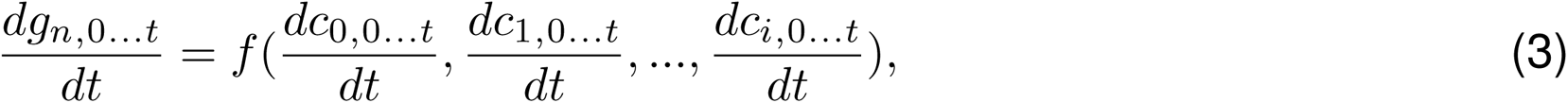

where *f* is a non-linear mapping function that can be solved by a neural network (NN) or random forest (RF). The former is particularly suitable to capture the non-linearity of the relationship between chromatin features and gene expression over time. Indeed, this architecture allows us to capture additional intricate patterns and relationships in the data compared to more classical predictive models such as the RF. The network encodes each chromatin derivative value, at every time point, into a high dimensional vector, and applies the Leaky-ReLU function. Then, the NN regresses each gene expression derivative, at each time point, as the output (Figure 5a). As mentioned before, we applied our modeling approach to genes that are associated with only one cCRE. However, this framework can be extended to cases where a gene is associated with multiple cCREs or where multiple chromatin features are considered, hence our model input is referred to as a tensor of chromatin *c* features at each time point *t*.

**Figure 5:**
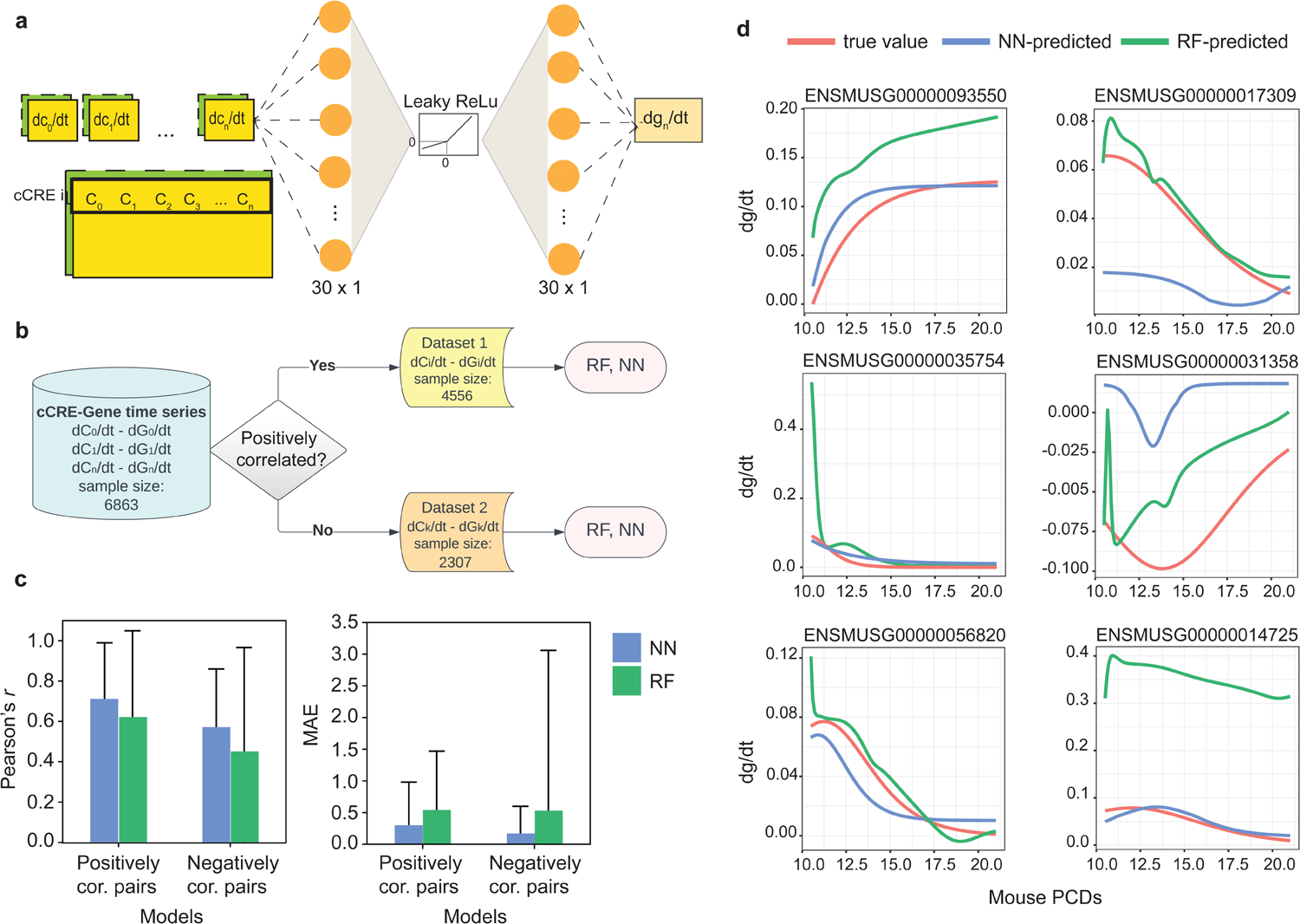
Modeling approach to predict changes in gene expression over time from changes in cCRE chromatin signals. **a:** Schematic representation of the neural network architecture. Each cCRE derivative, at each time point, is projected into a vector of 30×1 dimension followed by operating the Leaky ReLu function on each of its elements. Then, the gene expression derivative, at each time point, is predicted. **b:** Dataset construction for the two models. Each dataset consists of either negatively or positively cCRE-gene derivative pairs that are employed to train the Random Forest (RF) and Neural Network (NN) models. **c:** Performance evaluation of the models using Pearson’s r correlation coefficient (left panel) and Mean Absolute Error (MAE; right panel). Mean and standard deviation values were computed across all pairs in the test of each model. The NN model reported higher Pearson’s correlation and lower MAE compared to the RF. Standard deviations are larger in the RF compared to the NN model. **d:** Representative examples of predicted time-series gene expression derivatives by the NN (blue) and RF (green) models. True gene expression derivatives are shown in red.

Our training process strongly depends on the correlation direction between the time-series vectors of gene expression and chromatin derivatives. While the majority (66%) of cCRE-gene pairs showed positive correlation, a distinct subset displayed negative correlation (Supplementary Figure 4a), consistent with previous studies ^37–39^. Notably, this latter set of cCREs showed enrichment in motifs recognized by repressor TFs, such as ZBTB7A/B ^40–44^, suggesting a potential prototype for repressor-gene interactions. We thus divided the dataset into two subsets of positively and negatively correlated pairs (Figure 5b), and trained the network separately on each of the two subsets in a ratio of 80:20 for training and test sets.

Our predicted changes in gene expression showed overall positive correlation with the true changes in gene expression, as well as moderate-to-low mean absolute error (MAE; Figure 5c). Overall, the NN achieved higher performance than the RF model. Specifically, we reported a correlation of 0.71 ± 0.29 between the true and predicted values in the positively correlated pairs (average MAE 0.29 ± 0.69), and a correlation of 0.57 ± 0.29 in the negatively correlated pairs (average MAE = 0.16 ± 0.44). The RF model applied on the same task showed a correlation of 0.62 ± 0.43 with MAE of 0.53 ± 0.94 within the positively correlated pairs, and a correlation of 0.45 ± 0.50 with a MAE of 0.52 ± 2.54 for the negatively correlated pairs (Figure 5c). Overall, 99% and 100% of positively and negatively correlated cCRE-gene pairs, respectively, showed a positive correlation between true and NN-predicted gene expression derivatives, in contrast to the 89% and 83%, respectively, reported by the RF models (Supplementary Figure 4b). Representative examples of true and predicted changes in gene expression over time are shown in Figure 5d. Overall, these results demonstrate that chromatin accessibility changes within regulatory elements can be employed to accurately predict changes in the expression of putative target genes over time.

## Discussion

Time-series functional genomics assays offer a unique opportunity to investigate the transcriptional and chromatin kinetics of multiple genes and regulatory elements simultaneously. However, integrative analysis of these data requires a flexible framework that can uniformly model different types of signals without generating disparate parameter sets, thereby enabling direct comparison and biological interpretation of the inferred kinetics.

Here, we analyzed maps of chromatin accessibility generated during mouse brain development and identified cCREs with different types of accessibility kinetics among brain regions. Furthermore, we found that these kinetic patterns strongly differ between cCREs active in progenitors *vs.* more differentiated cell types. To analyze these patterns we adapted a well-known first-order differential equation–previously employed in other biological fields such as the study of bacterial population growth and decay–to model the kinetics of chromatin and gene expression changes over time. The ODE naturally accommodates some general principles of the kinetics governing these changes: first, that impulses of chromatin remodeling signals or transcriptional bursts manifest with rapid initial changes; second, that these changes eventually attain saturation over time due to biochemical constraints, such as the structure of histone complexes or the activation of feedback loop mechanisms. Previous methods have proposed log-like and exponential curves to model increases and decreases, respectively, of multi-omics signals ^21^. However, this follows the assumption that such changes begin in the early phases of the (latent) time course, and reach saturation by the end of the time course. Our framework instead relaxes these assumptions by incorporating a logistic curve, allowing more flexibility to capture diverse patterns of gene expression and chromatin signals.

To demonstrate the utility of this framework, we applied it to study the kinetics of chromatin accessibility during an eight-day time course of mouse development across three brain regions. We found that the majority of regulatory elements undergoing chromatin changes reached full accessibility or inaccessibility by the first post-natal day. Still, a fraction of these elements did not reach full accessibility by the end of the time course, especially those active in late-emerging cell types such as astrocytes and mature inhibitory neurons. Overall, this suggests that the kinetics of chromatin accessibility inferred in bulk recapitulate the emergence of cell type-specific patterns detected by single-cell experiments.

Our framework also allows us to investigate epigenome-transcriptome interactions without being constrained by predefined assumptions about their temporal dynamics. Specifically, we found that most of the cCRE-gene derivative pairs show a positive correlation, indicating that activator TFs may potentially bind to these cCREs. Conversely, cCREs showing negative correlation with their target genes may be recognized by repressor TFs. Based on this, we propose to independently model transcriptional and epigenetic changes using the same ODE and then to employ a neural network-based architecture to investigate their non-linear relationship over time. We specifically trained separate models for positively and negatively correlated cCRE-gene pairs that can capture activator-gene and repressor-gene interactions, respectively. This strategy is characterized by greater flexibility and allows us to model the kinetics of thousands of genes, compared to previous velocity methods which rely on a simplified view of gene regulation and can accurately fit only a restricted subset among thousands of genes ^21^. While we have applied this architecture to model the basic scenario of a gene regulated by a single cCRE, we anticipate that our approach to consider the time-series chromatin accessibility signal as a tensor can be scaled to more complex scenarios where multiple cCREs or chromatin features are employed to model changes in gene expression.

Although in the present study we have focused on modeling gene expression and chromatin accessibility data from bulk sequencing experiments, this framework is suitable to analyze other data modalities, and could also potentially accommodate time-resolved single cell measurements. We anticipate that applying *chronODE* to time-series data from various biological systems will help us to understand how alterations in transcriptional and epigenetic processes affect molecular pathways. These insights can be particularly valuable to identify potential drug targets and to understand their impact on cellular functionality across different tissues and cell types. As kinetic approaches begin to unveil molecular mechanisms underlying drug resistance in cancer ^45–47^, we anticipate that, in the long term, these kinetic maps of transcriptional and epigenetic processes will play a pivotal role in designing tailored therapeutic strategies.

## Author Contributions

B.B. and M.G. conceived the project. B.B., M.F., and M.G. designed the study. B.B., M.F., E.S.W., K.X., and S.X.L. performed the computational analyses. X.Y. contributed tools and ideas to perform computational analyses. B.B. and M.F. wrote the manuscript with the contribution of all authors.

## Supplementary Information

Supplementary Information is available for this paper.

## Methods

### Mouse brain developmental time-course

We analyzed maps of chromatin accessibility (DNase-seq and ATAC-seq) and gene expression (polyA+ RNA-seq) generated by the ENCODE consortium during eight time-points in three mouse fetal brain regions (forebrain, midbrain, and hindbrain) ^22,24^. DNase maps were available for postconception (PCD) days E10.5, E11.5, E14.5, and the first postnatal day (PN). ATAC-seq maps were available for PCDs E11.5, E12.5, E13.5, E14.5, E15.5, E16.5, and PN. RNA-seq data were available for all eight time points (PCD 10.5-PN) (Supplementary Table 1; https://www.encodeproject.org/matrix/type=Experiment&status=released&related_series.@type=OrganismDevelopmentSeries&replicates.library.biosample.organism.scientific_name=Mus+musculus&assay_title=ATACseq&life_stage_age=embryonic+10.5+days&life_stage_age=embryonic+11.5+days&life_stage_age=embryonic+12.5+days&life_stage_age=embryonic+13.5+days&life_stage_age=embryonic+14.5+days&life_stage_age=embryonic+15.5+days&life_stage_age=embryonic+16.5+days&life_stage_age=postnatal+0+days&biosample_ontology.term_name=forebrain&biosample_ontology.term_name=hindbrain&biosample_ontology.term_name=midbrain&assay_title=DNase-seq&assay_title=polyA+plus+RNA-seq). We built time-course matrices of signals for these two data modalities as we describe in the following sections.

### DNase- and ATAC-seq data processing

We downloaded the catalog of ENCODE ^25^ candidate *cis*-regulatory elements (cCREs) for the mouse genome from https://www.encodeproject.org/annotations/ENCSR412JPD/, which comprises 926,843 cCREs. We employed this catalog to construct a matrix of chromatin accessibility signals for each cCRE across the eight timepoints and the three brain regions. Given that ATAC-seq data were available from PCD E11.5 onward, in order to maximize the number of time points shared between the chromatin accessibility and gene expression maps, we integrated DNase-seq and ATAC-seq data in a single time course. Below we detail the steps of our DNase- and ATAC-seq data integration protocol.

#### Step 1: Identifying active cCREs during the time-course

For each of the three regions, we downloaded bigBed files of ATAC-seq pseudoreplicated narrow peaks available for each time point. We employed the BEDTools ^48^ (version 2.30.0) function `intersectBed` and identified 282,907 cCREs with ATAC-seq peaks in at least one time point. In the case of DNase-seq, pseudoreplicated peaks were unavailable. We therefore identified for each time point the peaks shared across all replicates using the BEDTools function `multiIntersectBed`. 316,549 cCREs reported DNase-seq peaks in at least one time point. We defined our set of 405,554 “active” cCREs as those that reported a DNase-seq and/or ATAC-seq peak. The list of bigBed files employed in this step is available in Supplementary Tables 4-5.

#### Step 2: Building a time-course matrix of chromatin accessibility for active cCREs

We downloaded ATAC-seq bigWig files (fold change over control; two replicates per time point and region; Supplementary Table 4) and computed the average signal in the cCRE window at each time point and replicate using the bigWigAverageOverBed tool. This yielded a 405,554 × 7 matrix for each replicate and region. We followed the same procedure for the DNase-seq signal (read-depth normalized signal; Supplementary Table 5), and obtained a 405,554 × 4 matrix for each replicate and region.

#### Step 3: Performing joint normalization and batch correction

We first performed joint quantile normalization on the ATAC- and DNase-seq signal matrices across replicates and time points using the R package preprocessCore ^49^. We then applied batch correction on the quantile-normalized matrices to remove unwanted effects due to experimental differences between the two assays. The input matrix consists of DNase-seq (PCD 10.5, 11.5, 14.5, PN; 2 replicates) and ATAC-seq (PCD 11.5, 12.5, 13.5, 14.5, 15.5, 16.5, PN; 2 replicates) signals. Shared time points 11.5, 14.5 and PN were employed to calibrate the differences between the two assays. Specifically, we first added a pseudocount of 1 to the signal matrix and performed a centered log-ratio transformation of the matrix with the R package mixOmics ^50^ (function `logratio.transfo`). After having the log-ratio normalized joint matrix of DNase and ATAC-Seq data, we conducted the batch effect correction with the R package Limma ^51^ (function `removeBatchEffect`), specifying the replicate and assay features as batch levels and the time course as design (Supplementary Figure 1).

### RNA-seq data processing

For each of the three brain regions, we downloaded from the ENCODE portal gene expression matrices of Transcript Per Million (TPM) values for the eight time points and the two biological replicates (mouse genome assembly version mm10, Gencode annotation version M21; Supplementary Table 6). We then normalized the data using center-log-ratio normalization and the limma function `removeBatchEffect`, specifying the replicate feature as batch level and the time course as design.

### Identifying dynamic cCREs and genes

To detect significant changes in cCRE chromatin accessibility over time, we used the R package maSigPro ^52^ with DNase- and ATAC-seq replicates handled internally. We conducted this analysis independently for each brain region. We employed the function `make.design.matrix()` to construct two different design matrices for linear (`degree = 1`) and quadratic (`degree = 2`) regression models. We then applied function `p.vector()` on each design matrix with the following parameters: `Q = 0.05`, `MT.adjust = “BH”`, `min.obs = 5`. For each region, we defined dynamic cCREs as those reporting a maSigPro FDR value *<* 0.01 in at least one of the two designs. We followed the same procedure to identify genes with significant expression changes over time, but first added a pseudocount of 10*^−^*^16^ to each gene expression value to avoid zero values.

### Identifying regulatory patterns of cCRE chromatin accessibility

We employed the R function `kmeans()` to group dynamic cCREs into clusters. Elbow plots indicated that, across the three regions, the most meaningful number of *k* -means clusters is 3, and that the three clusters correspond to downregulated (decreasing openness), early upregulated (openness increases early then stays high), and late upregulated (openness increases at the very end of gestation) cCREs (Figure 1b).

### The *chronODE* mathematical framework

We designed an ODE-based pipeline to capture trends in sparse time-series data. We employ equation (4) to describe the rate of change of chromatin accessibility or gene expression over time. The analytical solution of the ODE can be found in Supplementary Note 1. The pipeline, which we used to model chromatin accessibility of cCREs and RNA expression levels of genes, has two stages: linear interpolation and ODE fitting (Supplementary Figure 2a). The pipeline’s input takes the form of a two-dimensional matrix, with numeric time points as columns and elements (e.g. genes or cCREs) as rows. In our case, we used our eight time points in post-conception days. We chose to represent the postnatal time point as 21 PCD, since the standard length of a mouse pregnancy is typically in the range 19-21 days.

Since eight time points are insufficient to fit an ODE, the first step is data interpolation. We first used the function `linspace` (from the Python NumPy package) to generate a larger number of evenly spaced time points over the interval of the original time points. We chose to generate 105 time points. We then normalized the values in each row of the matrix to a range between zero and one, and created a new matrix with interpolated values for each of the new simulated time points using the Python Scipy package ^53^ (function `interp1d`).

Once we had a matrix of linearly interpolated values, we fitted an ordinary differential equation of the form

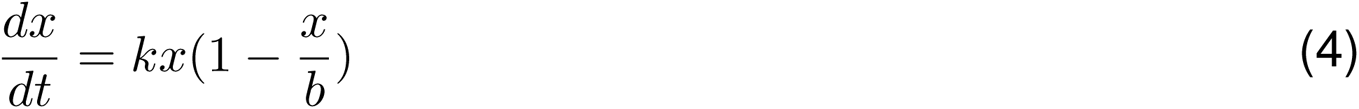

to each row. Given initial guesses for the *k* and *b* parameters (we chose 0.9 and 1.5, respectively), we then optimized the choice of *k* and *b* by fitting the equation (4) using the Scipy functions `curve fit` (maximum calling number equal to `5000`) and `odeint`, with the latter using LSODA, an adaptive steps algorithm ^54^. If the function cannot find optimized parameters in the predefined parameter space before hitting the maximum number of calls, it returns NAs. We then used the fitted parameters to model the values at the interpolated time points, and added a pseudocount of 10*^−^*^16^ to the first time point to avoid downstream divide-by-zero errors.

We fitted the ODE parameters using six sets of input: either a positive or negative initial guess for *k*, and three versions of the linearly interpolated data: the unshifted version, and versions raised or lowered by the maximum magnitude of the original normalized data. The fitted values were then shifted back by the same amount. Of these six fittings, we selected the set of parameters that yield fitted values with the lowest mean squared error compared to the linearly interpolated values. If all six fittings fail, the pipeline returns NAs for that row.

The pipeline outputs three matrices, with rows corresponding to the same cCRE/gene as the input matrix. The first two tables contain the fitted values and derivatives respectively, with a column for each of the interpolated time points. The third matrix contains the fitted parameters *k* and *b*, along with *T* = 1*/k*, MSE, and information about the vertical shift used to model each row.

With the output from the ODE pipeline, we first dropped the elements containing the NA (those whose trend cannot be captured by the ODE pipeline; Supplementary Figure 2a). We then did quality control by dropping the elements whose mean square error was among the highest 20%. With this filtered list, we clustered the cCREs based on their dynamic trend. We used a Convex-Concave approach to classify the dynamic trend. We identified the trend based on the fitted trajectory by connecting the fitted values’ first and last time points (Supplementary Figure 2b). After getting this decision line using the `linspace` function in the Numpy package (which has the same number of time points as the fitted trajectory), we calculated the difference between the fitted trajectory and the decision line. This difference is an array consisting of the difference at 105 time points. If the 105 difference values are all positive, we assign the cCRE with a “log-like” trend; if all the 105 difference values are negative, the cCRE is assigned to an “exponential” trend. If the signs of the 105 differences are a mixture of positive and negative, then the corresponding cCRE is believed to have the “logistic” trend.

### Identifying cCREs with region-specific kinetics

We employed the *k* parameter from the ODE pipeline output to identify cCREs with different kinetics across the three regions. As explained in the Results section “*chronODE* infers time-series trajectories and highlights cell type-specific kinetics of chromatin accessibility”, |k| summarizes the rate of change of a given cCRE or gene over time (Supplementary Figure 3a). We applied a normalization method and computed, for each cCRE, the ratio between |k| in a particular region and the average |k| across the three regions. We then performed *k* -means clustering separately on increasing and decreasing cCREs and identified in each case three main clusters, which correspond to cCREs reaching their highest relative rate of change in the forebrain, midbrain, and hindbrain (Figure 3a-b). For this analysis we considered only cCREs characterized by the same pattern (decreasing, early increasing and late increasing) across the three regions, and for visualization purposes we merged early and late increasing cCREs in one heatmap.

### Cell-type specific cCREs intersection analysis

We obtained sets of cell-type specific cCREs identified by single-cell (sc) ATAC-seq experiments performed in the mouse forebrain during the same time course as the bulk ATAC-seq experiments (PCD E11.5 through PN; specifically, Supplementary Table 4 from Preissl et *al.* ^23^). A description of the main cell types corresponding to each cCRE cell-type specific set can be found in Supplementary Table 3. We assigned each of our forebrain dynamic cCREs to a specific cell-type specific set by employing the BEDTools function `intersectBed`.

### Linking cCREs to putative target genes

We designated a target gene for each cCRE based on linear distance. We used the BEDTtools `closest` utility for this calculation. This methodology routinely assigns many cCREs to a single target gene. A cCRE, however, can only be assigned multiple targets if two or more genes are tied for “closest”, usually because they all overlap the cCRE. BEDTools `closest` assigns a distance of 0 to all overlaps. Having linked genes to dynamic cCREs, each of which had already been assigned a regulatory pattern (see above), we used the UpSet R package to visualize how many genes are linked to every possible combination of cCRE regulatory patterns (Figure 4a). We then divided the genes by the number of cCRE patterns that target them.

### Properties of mono- *vs.* multi-pattern genes

To examine the properties of genes targeted by different numbers of patterns, we performed Gene Ontology (GO) analysis and transcription-factor (TF) motif enrichment analysis on the genes linked to one, two, and three regulatory patterns from each brain region. For the GO analysis, we used the GOStats R library ^55^. As universe gene set for the GO enrichment analysis we used the set of protein-coding genes from the Gencode M21 mouse annotation and a false-discovery rate cutoff of FDR *<* 0.01 to identify significantly enriched GO terms related to biological processes. We also used the Homer motif discovery tool ^56^ to find TF motifs that were significantly enriched in each group of genes. We used the `findMotifs.pl` script and HOMER’s built-in mouse promoter set to identify TF motifs that are significantly enriched in the promoter region of each set of genes.

### Motif analysis of correlation groups

In order to decipher the difference between the set of cCREs with positive versus negative correlations, we conducted TF motif analysis. We downloaded the reference fasta file from ENCODE and used the BEDTools `getfasta` command to generate the fasta files of the two sets of cCREs. We then used STREME ^57^ to discover ungapped motifs that are relatively enriched in each of the two sets of cCREs using the other set as control sequences. In this process we chose the patience to be 10, so the software would stop searching for motifs when ten consecutive non significant motifs have been found. Then we compared the motifs we discovered against the database HOCOMOCO (version 11).

### Predicting changes in gene expression from changes in chromatin accessibility Neural Network Model

In order to predict the gene expression derivatives over time we used the chromatin accessibility derivatives as inputs for a non-linear neural network with the following architecture: Num features = 1, Degree

`= 30`, `Linear(num features, degree, bias=True)`, `LeakyReLU(0.4,inplace=True)`, and `Linear(degree, 1)`. Specifically, each of the chromatin derivatives, at each time point, serves as an input for the network (Figure 5a). The input is defined as a 3D tensor where the first position stands for the number of samples, the second position is the number of cCRE-related-features (which in this paper is 1), and the third position is a 105 dimensional vector of the different epigenetic signals over time. The output is also a vector of a 3D tensor, where the first position points to the same number of samples as the inputs, each corresponding to one gene-related derivative across the 105 time points. We used PyTorch ^58^ to train the network and set the batch size to 4 along with a mean absolute error as the loss function and a learning rate of 0.001 using the Adam optimizer. Finally, the network was trained with 3000 epochs. We combined the three regions for the purposes of this model, but split the gene-cCRE pairs into two groups depending on whether the chromatin accessibility derivatives were positively or negatively correlated with the gene expression derivatives. We trained and tested the neural network separately on these two groups in a ratio 80:20.

### Random Forest Model

We also used a random-forest-based regression model to predict gene expression derivatives based on chromatin openness derivatives. To ensure easily comparable results, we used the same 105-timepoint input and output matrix formats for the random forest model as for the neural network described above, and used the same correlation-based split. Thus, the 80:20 train and test sets were identical to the ones employed by the Neural Network. We used the scikit-learn ^59^ function `RandomForestRegressor` with 100 trees and a default depth of 2.

### Quantification and statistical analysis

All statistical analyses were performed using the R or Python languages, as specified in the Methods and/or figure legends. Unless otherwise specified, plots were made with the R package ggplot2^60^ or the Python package matplotlib ^61^. All box plots depict the first and third quartiles as the lower and upper bounds of the box, with a band inside the box showing the median value and whiskers representing 1.5x the interquartile range.

