## Supplementary Information for "*chronODE*: A framework to integrate time-series multi-omics data based on ordinary differential equations combined with machine learning"

Supplementary Figures 1-4
Supplementary Tables 1-6
Supplementary Note

**Supplementary Figure 1**

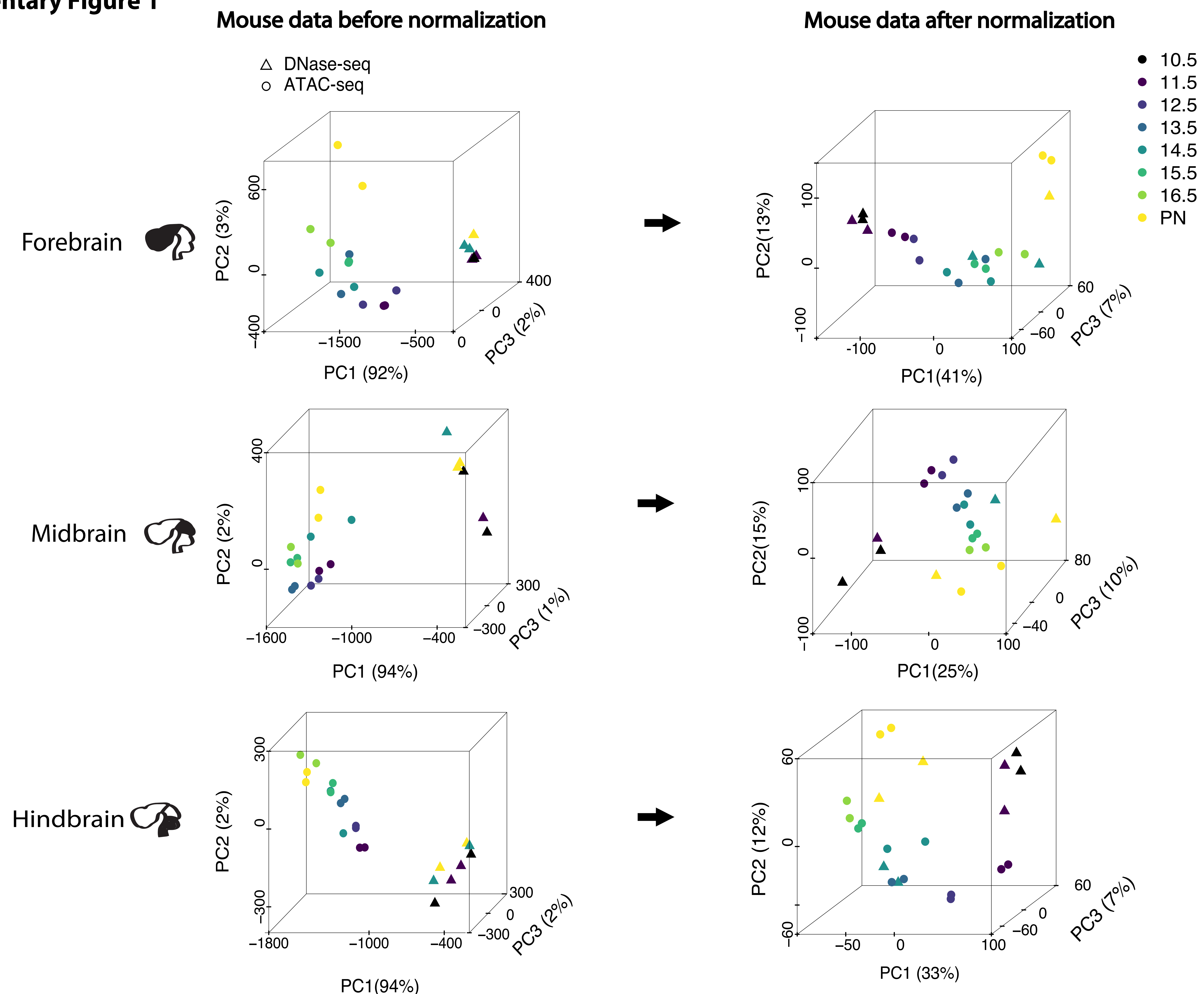

**Supplementary Figure 1: Data normalization protocol to integrate time-series DNase-seq and ATAC-seq data into a single time course.** Principal Component Analysis (PCA) of DNase- and ATAC-seq signals before (left) and after (right) applying data normalization. Each observation in the PCA space corresponds to a time point and assay (time points are color-coded; DNase-seq: triangle points; ATAC-seq: circle points). We employed the time-series chromatin signal at the 405,554 active cCREs as variables for the PCA. Following our protocol (see Methods section "DNase- and ATAC-seq data processing"), we could observe more consistent integration of the time points profiled by the two assays. The percentage of variance explained by each PC is shown between parenthesis.

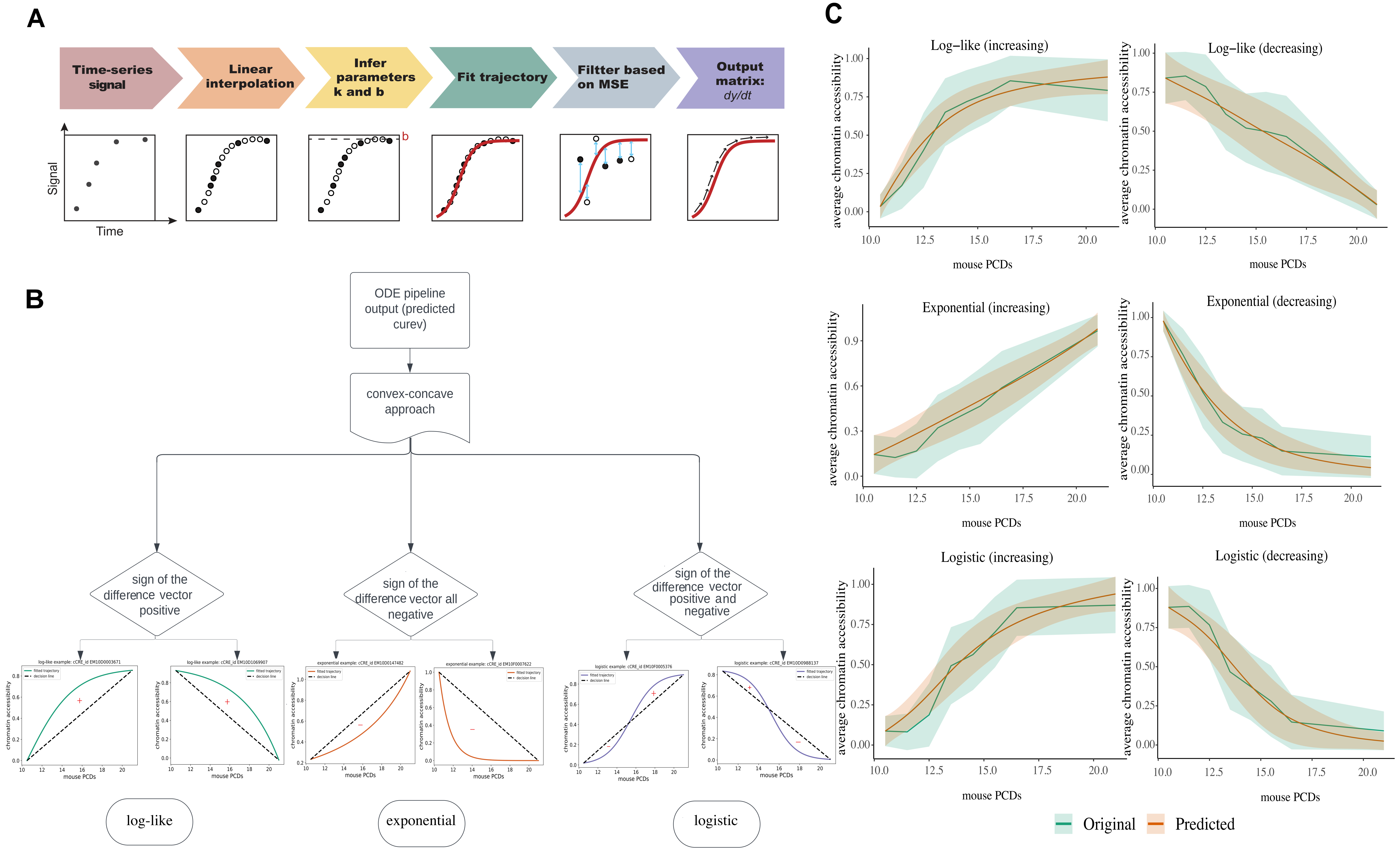
A
Time-series signal
Linear interpolation
Infer parameters k and b
Fit trajectory
Filter based on MSE
Output matrix: dy/dt
Signal
Time
b
B
ODE pipeline output (predicted curev)
convex-concave approach
sign of the difference vector positive
sign of the difference vector all negative
sign of the difference vector positive and negative
log-like
exponential
logistic
C
Log−like (increasing)
Log−like (decreasing)
Exponential (increasing)
Exponential (decreasing)
Logistic (increasing)
Logistic (decreasing)
average chromatin accessibility
mouse PCDs
Original
Predicted

**Supplementary Figure 2: Overview of the chronODE modeling framework.**

**a**. Schematic showing the major steps of the pipeline (see Methods section “The chronODE mathematical framework”). chronODE takes as input the original time-series signal for a given gene or cCRE and performs a linear interpolation to increase the number of time points with available signal. It then fits equation (1) on the interpolated data to compute parameters k and b and the time-series derivatives of the signal trajectory. It then computes the Mean Squared Error (MSE) between the fitted trajectory and the original data and discards poorly fitted cases. The main output is the matrix of time-series derivatives. **b.** Schematic showing the classification of time-series trajectories following the convex-concave approach. Log-like trajectories are defined by a vector of positive residuals with respect to the regression line fitting the lower and upper quantiles. Exponential trajectories are defined by a vector of negative residuals. Logistic trajectories are defined by a mixture of positive and negative residuals. **c.** Average chromatin accessibility trajectories of increasing (left) and decreasing (right) cCREs modeled by chronODE for different curve types (exponential, log-like, logistic). For each group we show both averaged observed (green) and predicted (orange) trajectories.

**Supplementary Figure 3**

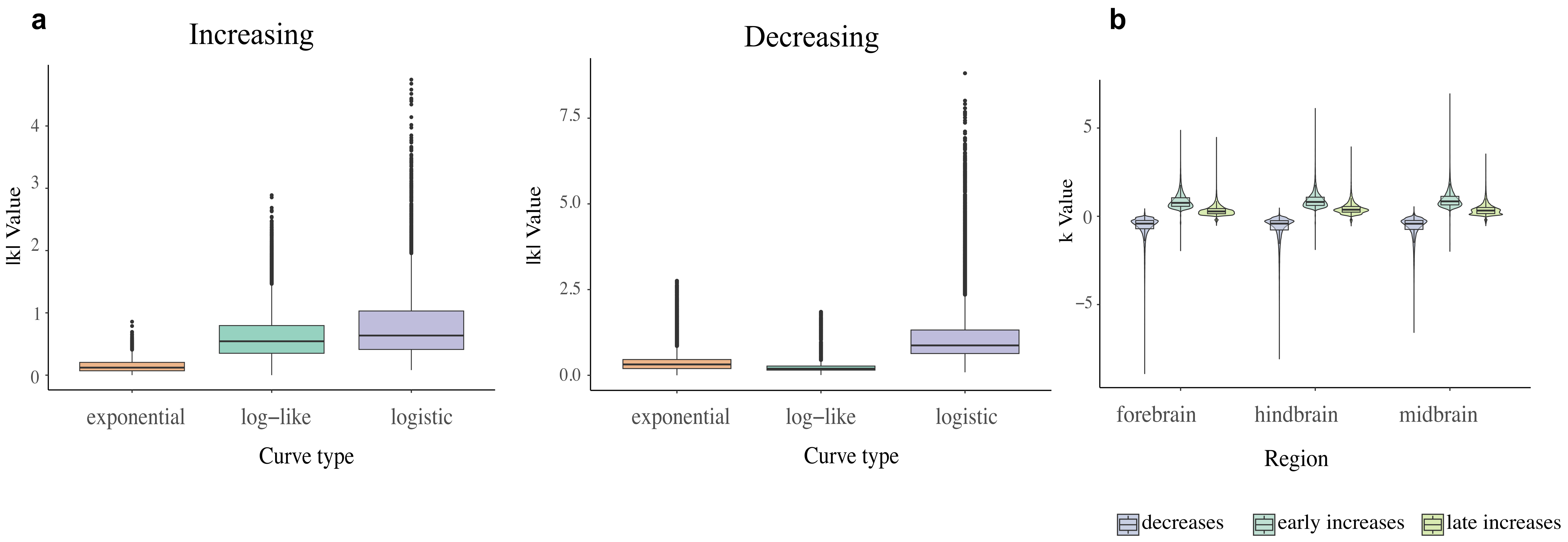

**Supplementary Figure 3: Interpreting the biological meaning of *chron*ODE kinetic parameters.**

**a:** box plots showing the distribution of $|k|$ values (y axis) across different curve types (exponential, log-like, logistic; x axis) in increasing (left) and decreasing (right) chromatin accessibility patterns. **b:** Box plot showing the distribution of $k$ values (y axis) for different chromatin accessibility patterns (decreases, early increases, late increases; color-coded) across brain regions (x axis).

### Supplementary Figure 4

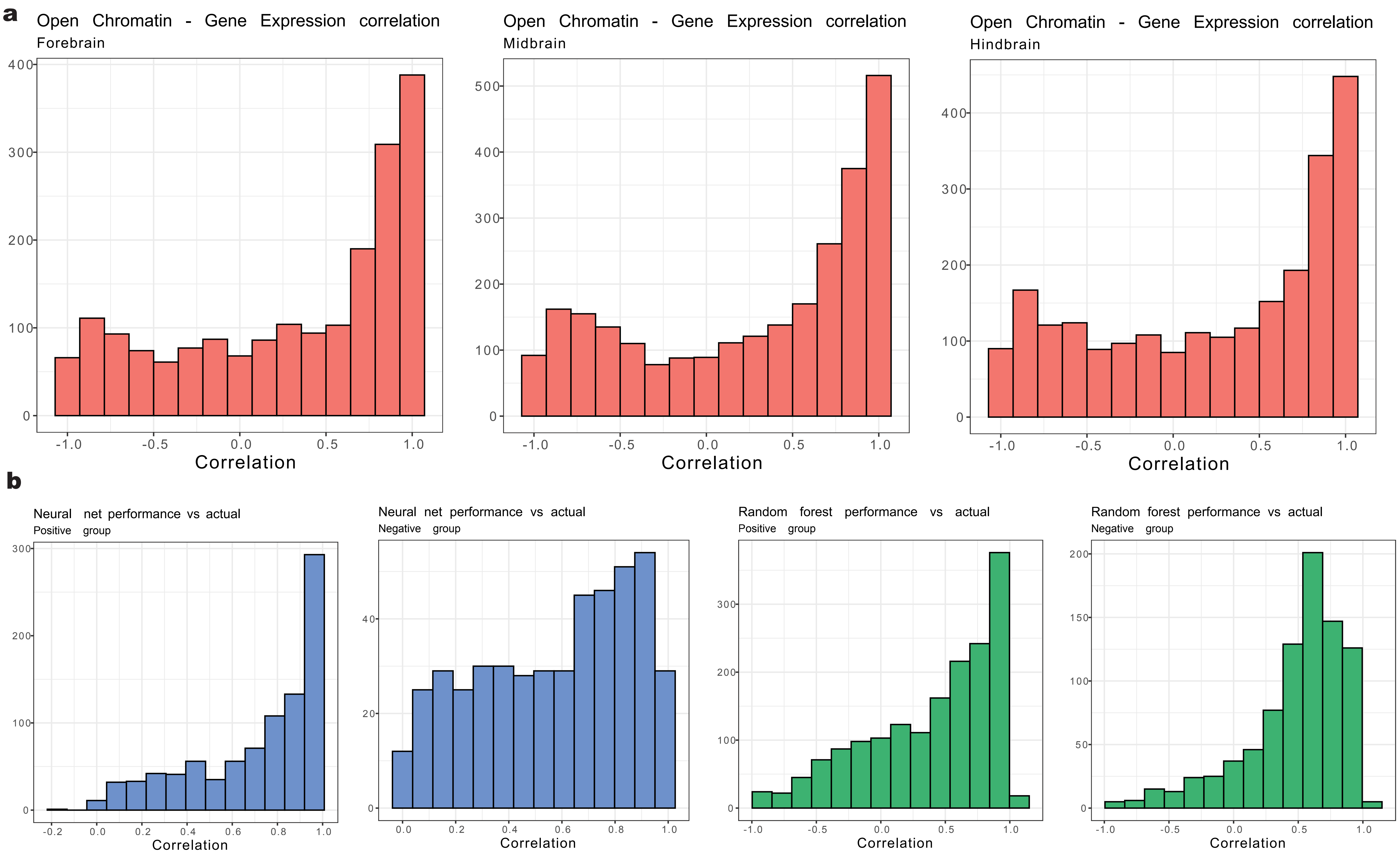

**Supplementary Figure 4: Correlation of derivatives of gene expression with chromatin accessibility and model predictions.**
**a:** Distribution of the correlation between the derivative of chromatin accessibility of a cCRE and the derivative of RNA expression level at its linked gene, for one cCRE-one gene pairs in the three brain regions. **b:** Distribution of the correlation between actual and predicted gene expression derivatives, for neural net (blue) and random forest (green) models. Gene-cCRE pairs were split into positive and negative groups based on the sign of the actual correlation between chromatin accessibility and gene expression.

| mouse PCDs | eq. human PCWs with heterochronies | eq. human PCWs without heterochronies |
|---|---|---|
| 10.5 | 4-6 | 4-6 |
| 11.5 | 7-9 | 4-6 |
| 12.5 | 7-9 | 7-9 |
| 13.5 | 10-12 | 7-9 |
| 14.5 | 10-12 | 7-9 |
| 15.5 | 13-15 | 10-12 |
| 16.5 | 16-18 | 10-12 |
| PN | 25-27 | 19-21 |

**Supplementary Table 1**. Comparison of mouse post-conception days with human post-conception weeks with and without heterochronies. Equivalence between mouse and human time courses was obtained from `https://www.translatingtime.org/translate/` (Workman *et al.*, 2013).

| Pattern | Type of change | Forebrain (%) | Midbrain (%) | Hindbrain (%) |
|---|---|---|---|---|
| Decreases | Concordant | 98 | 99 | 98 |
| | Discordant | 2 | 1 | 2 |
| Early increases | Concordant | 51 | 63 | 50 |
| | Discordant | 49 | 37 | 50 |
| Late increases | Concordant | 59 | 48 | 61 |
| | Discordant | 41 | 52 | 39 |

**Supplementary Table 2**. Proportion (%) of cCREs in each category (decreases, early increases, and late increases) that undergo concordant or discordant changes among the three brain regions.

| Cluster *k* | Label | Cell type annotation |
|---|---|---|
| 1 | RG1-2 | Radial Glia 1-2 |
| 2 | eAC | Astrocytes |
| 3 | ubiq. 1 | Radial Glia 1-4, Astrocytes |
| 4 | ubiq. 2 | Radial Glia 4, Astrocytes, Excit. Neurons 1-2 |
| 5 | ubiq. 3 | Radial Glia 1-4, Astrocytes, Inhib. Neurons 2, Excit. Neurons 1 |
| 6 | ubiq. 4 | All cell types but Excit. Neurons 2 and Erythromyeloid Progenitors |
| 7 | ubiq. 5 | All cell types but Radial Glia 1-2,4 and Erythromyeloid Progenitors |
| 8 | eIN3 | Inhibitory Neurons 3 |
| 9 | eIN2-4 | Inhibitory Neurons 2-4 |
| 10 | RG3, eIN2 | Radial Glia 3 and Inhibitory Neurons 2 |
| 11 | eIN4 | Inhibitory Neurons 4 |
| 12 | eEX1 | Excitatory Neurons 1 |
| 13 | eEX2 | Excitatory Neurons 2 |
| 14 | EMP | Erythromyeloid Progenitors |
| 15 | ubiq. 6 | All cell types |

**Supplementary Table 3**. Description of cell-type specific peaks identified by time-series single-cell ATAC-seq experiments performed in the mouse forebrain. The cell type annotations follow Figure 4 of Preissl *et al.,* 2018.

| Assay | Experiment ID | bigBed file ID | bigWig file ID | Region | Time point |
|---|---|---|---|---|---|
| ATAC | ENCSR211OCS | ENCFF750OEN | ENCFF116OYR, ENCFF231JIR | midbrain | 0 days |
| ATAC | ENCSR310MLB | ENCFF178FIA | ENCFF171GQJ, ENCFF374YJI | forebrain | 0 days |
| ATAC | ENCSR312LQX | ENCFF855NTE | ENCFF981VLS, ENCFF752FVM | hindbrain | 0 days |
| ATAC | ENCSR012YAB | ENCFF693MQK | ENCFF067IQO, ENCFF129DMB | hindbrain | 11.5 days |
| ATAC | ENCSR273UFV | ENCFF582AQM | ENCFF609FLS, ENCFF014TVL | forebrain | 11.5 days |
| ATAC | ENCSR382RUC | ENCFF697ALQ | ENCFF298FKT, ENCFF886SFL | midbrain | 11.5 days |
| ATAC | ENCSR088UYE | ENCFF146WID | ENCFF533PFX, ENCFF642QRZ | hindbrain | 12.5 days |
| ATAC | ENCSR154BXN | ENCFF818SNO | ENCFF460PDT, ENCFF860MZS | midbrain | 12.5 days |
| ATAC | ENCSR559FAJ | ENCFF724CPC | ENCFF314LIV, ENCFF586FIG | forebrain | 12.5 days |
| ATAC | ENCSR176BYZ | ENCFF894TOH | ENCFF953AOC, ENCFF280BOI | hindbrain | 13.5 days |
| ATAC | ENCSR819QOJ | ENCFF490UUC | ENCFF063FOL, ENCFF355SPQ | midbrain | 13.5 days |
| ATAC | ENCSR903GMO | ENCFF761PYJ | ENCFF278ROE, ENCFF161RRE | forebrain | 13.5 days |
| ATAC | ENCSR384JBF | ENCFF019TUF | ENCFF927RDV, ENCFF518JWO | midbrain | 14.5 days |
| ATAC | ENCSR798FDL | ENCFF859DRE | ENCFF572TQP, ENCFF823LHI | hindbrain | 14.5 days |
| ATAC | ENCSR810HQR | ENCFF197DQD | ENCFF092VLN, ENCFF039BNT | forebrain | 14.5 days |
| ATAC | ENCSR468GUI | ENCFF821IXB | ENCFF356BYP, ENCFF106YOY | midbrain | 15.5 days |
| ATAC | ENCSR662KNY | ENCFF748XVF | ENCFF566SYW, ENCFF985BES | hindbrain | 15.5 days |
| ATAC | ENCSR976LWP | ENCFF501WGN | ENCFF398CBN, ENCFF764JRZ | forebrain | 15.5 days |
| ATAC | ENCSR096JCC | ENCFF755XMC | ENCFF662ETT, ENCFF074YXW | midbrain | 16.5 days |
| ATAC | ENCSR623GSD | ENCFF586TSG | ENCFF770SZX, ENCFF800APZ | hindbrain | 16.5 days |
| ATAC | ENCSR836PUC | ENCFF779DKA | ENCFF365KJF, ENCFF807OVL | forebrain | 16.5 days |

**Supplementary Table 4**. Experiment and file identifiers for ATAC-seq experiments obtained from the ENCODE portal `https://www.encodeproject.org/`. Identifiers for DNase-seq experiments are provided in the following page (Supplementary Table 5).

| Assay | Experiment ID | bigBed file ID | bigWig file ID | Region | Time point |
|---|---|---|---|---|---|
| DNase | ENCSR469VGZ | ENCFF868GJZ, ENCFF233WJN, ENCFF314FJW, ENCFF500TCE | ENCFF822FDB, ENCFF286LUT | hindbrain | 0 days |
| DNase | ENCSR767AJS | ENCFF305HSJ, ENCFF125JIF, ENCFF277HPA, ENCFF650WJJ | ENCFF325FZS, ENCFF584BTI | midbrain | 0 days |
| DNase | ENCSR791AJY | ENCFF217LRD, ENCFF928NJG | ENCFF727CYI | forebrain | 0 days |
| DNase | ENCSR289BTM | ENCFF096PEE, ENCFF020ZKO, ENCFF862MJH, ENCFF920TRY | ENCFF790CVH, ENCFF072IWN | hindbrain | 10.5 days |
| DNase | ENCSR756SPS | ENCFF880ICH, ENCFF316KAS, ENCFF287EYQ, ENCFF544FGH | ENCFF506ZSC, ENCFF044ISO | forebrain | 10.5 days |
| DNase | ENCSR773SAG | ENCFF181QNC, ENCFF317OFK, ENCFF534WLB, ENCFF321PTE | ENCFF293YCS, ENCFF330OMV | midbrain | 10.5 days |
| DNase | ENCSR014SFF | ENCFF050WLI, ENCFF993LQC, ENCFF779DJK, ENCFF271HGS | ENCFF639LUT, ENCFF580FOS | forebrain | 11.5 days |
| DNase | ENCSR292QBA | ENCFF382EUY, ENCFF524WVA | ENCFF414SMG | midbrain | 11.5 days |
| DNase | ENCSR358ESL | ENCFF113JBH, ENCFF656GBX, ENCFF465HTG, ENCFF344ZDH | ENCFF587MWA, ENCFF609ATS | hindbrain | 11.5 days |
| DNase | ENCSR179PIH | ENCFF702YOI, ENCFF083VZJ, ENCFF858OHC, ENCFF965UXS | ENCFF274NPS, ENCFF537XKO | hindbrain | 14.5 days |
| DNase | ENCSR337EDG | ENCFF821UVA, ENCFF854HCQ, ENCFF225UEN, ENCFF093PQD | ENCFF237YNH, ENCFF931CQE | forebrain | 14.5 days |
| DNase | ENCSR367FCW | ENCFF310VOE, ENCFF876PHQ | ENCFF539CBL | midbrain | 14.5 days |

| Assay | Experiment ID | Quantification tsv file ID | Region |
|---|---|---|---|
| RNAseq | ENCSR017JEG | ENCFF892WXB, ENCFF851KEG | hindbrain |
| RNAseq | ENCSR080EVZ | ENCFF719ADL, ENCFF029UVS | forebrain |
| RNAseq | ENCSR160IIN | ENCFF696BTU, ENCFF042VCB | forebrain |
| RNAseq | ENCSR185LWM | ENCFF270DCV, ENCFF565AWX | forebrain |
| RNAseq | ENCSR285WZV | ENCFF310GSV, ENCFF858ZON | hindbrain |
| RNAseq | ENCSR304RDL | ENCFF476ADM, ENCFF145PTV | forebrain |
| RNAseq | ENCSR307BCA | ENCFF094JLI, ENCFF184FWR | midbrain |
| RNAseq | ENCSR343YLB | ENCFF743IEH, ENCFF954BEO | midbrain |
| RNAseq | ENCSR362AIZ | ENCFF484FBW, ENCFF143OBR | forebrain |
| RNAseq | ENCSR367ZPZ | ENCFF238YUA, ENCFF762ZJZ | midbrain |
| RNAseq | ENCSR401BSG | ENCFF195CMT, ENCFF804NPY | hindbrain |
| RNAseq | ENCSR420QTO | ENCFF928MQD, ENCFF046RSQ | hindbrain |
| RNAseq | ENCSR557RMA | ENCFF670AQP, ENCFF624EQM | midbrain |
| RNAseq | ENCSR559TRB | ENCFF304ILZ, ENCFF876LKY | hindbrain |
| RNAseq | ENCSR647QBV | ENCFF698XIB, ENCFF649WEQ | forebrain |
| RNAseq | ENCSR719NAJ | ENCFF492ODZ, ENCFF532MRQ | midbrain |
| RNAseq | ENCSR752RGN | ENCFF080PBH, ENCFF890FJF | forebrain |
| RNAseq | ENCSR760TOE | ENCFF606UHO, ENCFF434CSI | hindbrain |
| RNAseq | ENCSR764OPZ | ENCFF875HME, ENCFF126VCW | midbrain |
| RNAseq | ENCSR792RJV | ENCFF867NEL, ENCFF189APQ | midbrain |
| RNAseq | ENCSR908JWT | ENCFF521YOL, ENCFF840AXS | midbrain |
| RNAseq | ENCSR921PRX | ENCFF960KJV, ENCFF356CTG | hindbrain |
| RNAseq | ENCSR943LKA | ENCFF169CJH, ENCFF997ZPU | hindbrain |
| RNAseq | ENCSR970EWM | ENCFF794PWS, ENCFF088OEQ | forebrain |

**Supplementary Table** . 6. Experiment and file identifiers for RNA-seq experiments obtained from the ENCODE portal https://www.encodeproject.org/.

### Supplementary Note 1

#### ANALYTICAL SOLUTION OF THE POPULATION DYNAMIC ODE

**Theorem.** *Given*

$$\frac{dy}{dt} = ky(1 - \frac{y}{b})$$

*and the initial point* $(t_0, y_0)$
*we have:*

$$|y(t)| = |\frac{bCe^{kt}}{b + Ce^{kt}}| \tag{1}$$

*where* $C = e^c$

$$c = ln|y_0| - ln|1 - \frac{y_0}{b}| - kt_0$$

Proof:

$$\frac{dy}{dt} = ky(1 - \frac{y}{b})$$

$$\frac{dy}{y(1 - \frac{y}{b})} = kdt$$

$$(\frac{1}{y} + \frac{\frac{1}{b}}{(1 - \frac{y}{b})})dy = kdt$$

$$ln|y| - ln|1 - \frac{y}{b}| = kt + c$$

Then, we have:

$$ln|y_0| - ln|1 - \frac{y_0}{b}| = kt_0 + c$$

$$c = ln|y_0| - ln|1 - \frac{y_0}{b}| - kt_0$$

$$ln|y| - ln|1 - \frac{y}{b}| = kt + c$$

$$\frac{|y|}{|1 - \frac{y}{b}|} = Ce^{kt}$$

$$|y| = |\frac{bCe^{kt}}{b + Ce^{kt}}|$$

where $C = e^c$

**Theorem.** *Given*

$$y(t) = \frac{bCe^{kt}}{b + Ce^{kt}}$$

*we have:*

$$\lim_{t\to\infty} y(t) = \begin{cases} 0 & \text{if } k < 0 \\ b & \text{if } k > 0 \end{cases}$$

Proof:
Notice:

$$\begin{aligned} \frac{1}{y(t)} &= \frac{b + Ce^{kt}}{bCe^{kt}} \\ &= \frac{1}{Ce^{kt}} + \frac{1}{b} \end{aligned}$$

When $k > 0$, we have

$$Ce^{kt} \to \infty$$

when $t \to \infty$
So

$$\lim_{t\to\infty} \frac{1}{y(t)} = \frac{1}{b}$$
$$\lim_{t\to\infty} y(t) = b$$

When $k < 0$, we have

$$Ce^{kt} \to 0$$

when $t \to \infty$
So

$$\lim_{t\to\infty} \frac{1}{y(t)} = \infty$$
$$\lim_{t\to\infty} y(t) = 0$$
